# Fecal microbiota transplantation efficacy for *Clostridium difficile* infection: A trans-kingdom battle between donor and recipient gut microbiomes

**DOI:** 10.1101/2020.05.27.120386

**Authors:** Negin Kazemian, Milad Ramezankhani, Aarushi Sehgal, Faizan Muhammad Khalid, Amir Hossein Zeinali Kalkhoran, Apurva Narayan, Gane Ka-Shu Wong, Dina Kao, Sepideh Pakpour

## Abstract

Fundamental restoration ecology and community ecology theories can help us better understand the underlying mechanisms of fecal microbiota transplantation (FMT) and to better design future microbial therapeutics for recurrent *Clostridium difficile* infections (rCDI) and other dysbiosis-related conditions. In a single cohort study, stool samples were collected from donors and rCDI patients one week prior to FMT (pre-FMT) as well as from patients one week following FMT (post-FMT). Using metagenomic sequencing and machine learning methods, our results suggested that the FMT outcome is not only dependent on the ecological structure of the recipients, but also the interactions between the donor and recipient microbiomes, both at the taxonomical and functional levels. Importantly, we observed that the presence of specific bacteria in donors (*Clostridiodes* spp., *Desulfovibrio* spp., *Odoribacter* spp. and *Oscillibacter* spp.) and the absence of specific fungi (*Yarrowia* spp.) and bacteria (*Wigglesworthia* spp.) in recipients prior to FMT could accurately predict FMT success. Our results also suggested a series of interlocked mechanisms for FMT success, including the repair of the disturbed gut microbial ecosystem by transient colonization of nexus species followed by secondary succession of bile acid metabolizers, sporulators, and short chain fatty acid producers. Therefore, a better understanding of such mechanisms can be fundamental key elements to develop adaptive, personalized microbial-based strategies for the restoration of the gut ecosystem.

**Importance:** There have been a number of studies focusing on understanding the underlying mechanisms in FMT treatment, which can accordingly be used for the optimization of future treatments. However, the current scientific lens has mainly had a uni-kingdom major focus on bacteria, leading to the proposition of the existence of FMT “super-donors”. On the contrary, our preliminary study here suggests that FMT is not necessarily a ‘one stool fits all’ approach and that donor-recipient cross-kingdom microbiota interactions, along with their short-term fluctuations in the gut, bring profound implications in FMT success. The results also conceptualize a series of interlocked mechanisms for FMT success, including first repairing the disturbed gut microbial ecosystem by transient species, followed by secondary succession of indigenous or exogenous bile acid metabolizers, sporulators, and short chain fatty acid producers.

## Introduction

Antibiotics are the primary treatment method for *Clostridium difficile* infections (CDI); however, the negative impact on the diversity, composition, and functionality of gut microbiota results in recurrent CDI (rCDI) (1, 2) requiring fecal microbiota transplantation (FMT). FMT is a strategy for the restoration of a disturbed microbial ecosystem and reinstatement of lost microbial functional networks. Although highly effective in the treatment of rCDI as well as promising in several other diseases (2–8), FMT carries infectious and non-infectious risks (9–12). In addition, under each specific disease scenario, it is crucial to understand how microbial ecosystems reassemble overtime after FMT and which microbial strains are the determining factors in this dynamic process. For rCDI treatment, the current scientific lens has mainly had a uni-kingdom major focus on bacteria. It has been suggested that an ideal donor should have high *Lachnospiraceae* and *Ruminococcaceae* (13), which are also positively associated with secondary bile acids that inhibit CDI germination (1). Increased *Clostridium Scindens* in donors has also shown a positive correlation with FMT efficacy and outcomes via the production of secondary bile acids (14). Moreover, FMT restores short chain fatty acids (SCFAs) metabolism, with immune modulatory effects in rCDI patients (15). SCFAs and butyrate producing bacteria have been found to decrease the induction of proinflammatory cytokines and promote the differentiation of colonic Treg cells, leading to the attenuation of colitis in mice and humans (16, 17). In addition, anaerobic, endospore-forming Firmicutes are dominant members of gut microbiota that can produce SCFAs (18), which allow organisms to enter metabolically dormant states that aid in their survival and transmission to new hosts (19). Thus, the oral delivery of SER-109, composed of sporulating bacteria, remains a promising therapeutic approach for rCDI treatment (20, 21). Furthermore, a critical consideration for FMT efficacy and durability is that the microbial consortium of the donors is not the only key player. The existing endogenous microbiome in recipients can also play a significant role in determining the colonization of those exogenous species. For example, focusing on bacterial engraftment, Smillie et al. (22) suggested that selective forces in the patient’s gut (host control), rather than input dose dependence (bacterial abundance in the donor and patient), determines bacterial abundance after FMT and, subsequently, its efficacy. In contrast, a number of studies suggest that FMT success is only dependent on the bacterial diversity and composition of the stool donor, leading to the proposition of the existence of FMT super-donors (3, 23).

Beyond the gut bacterium, few studies have examined the role of gut mycobiome and virome on FMT efficacy. For example, Zuo and colleagues found a negative relationship between the abundance of fungi such as *Candida albicans* in donor stool and FMT efficacy (24). Over the last decade, phages have gained increasing attention for therapeutic use due to their specificity (25). The reduction in the abundance of *Caudovirales* bacteriophages and an increase in *Microviridae* abundance, specifically higher abundance of *Eel River basin pequenovirus* as a potential Proteobacteria predator, were shown to be related to FMT efficacy in CDI patients (26, 27). Using targeted refined phage therapy, Nale et al. (28) used a cocktail of four *C. difficile* Myoviruses (CDHM1, 2, 5, and 6) to eradicate the CDI in a batch fermentation model, which suggests that a combination of bacteriophages may be needed to treat CDI. More recently, rCDI in five patients was prevented using sterile fecal filtrate, void of live bacteria (29). Contrary to these, a study by Meader et al. (30) showed that bacteriophages alone weren’t sufficient to eradicate CDI. These studies emphasize that in order to uncover mechanisms involved in FMT efficacy, it is fundamental to include the relative contribution of all domains and consider the microbiome-associated ecosystem heterogeneity in both donors and recipients. To this end, we specifically investigated whether FMT super-donors exists for rCDI treatment, or whether the donor-recipient compatibility and short-term fluctuations in the gut microbiomes (a combination of bacteria, fungi, archaea, and viruses) of both donors and recipients have profound implications in FMT success.

## Results

Among recipients, 9/17 patients were successfully treated with a single FMT (53% successful FMT), while 8 patients failed the first FMT and required a second procedure. There was no difference between the two groups in factors of age, sex, or duration of CDI (31).

We found no significant difference in alpha diversities of different organisms in stool samples provided by donors used for all patients whether the treatment outcome was successful or not (Kruskal-Wallis test, p>0.05, Fig. 1A-E). For the recipients, no significant differences in alpha diversities were observed between successful and failed pre-FMT samples (Kruskal-Wallis test, Fig. 1). There was a significant increase in the bacterial (Fig. 1A) and fungal (Fig. 1C) alpha diversities (Shannon diversity index) in post-FMT stool samples after successful FMT (Wilcoxon test, adjusted p-value<0.0001 and p-value=0.002, respectively), but not failed ones. No significant changes in this index were seen post-FMT in archaeal, protozoan, and viral diversities (Fig. 1B, D, E). It is important to note that although no significant changes were observed for viral diversities, the interplay between phages and bacteria in rCDI and FMT treatment is intricate especially due to technical limitations of viral enrichment in biological samples, extraction and sequencing library bias towards dsDNA viruses, and removal of ssDNA and RNA viruses.

**Fig 1.**
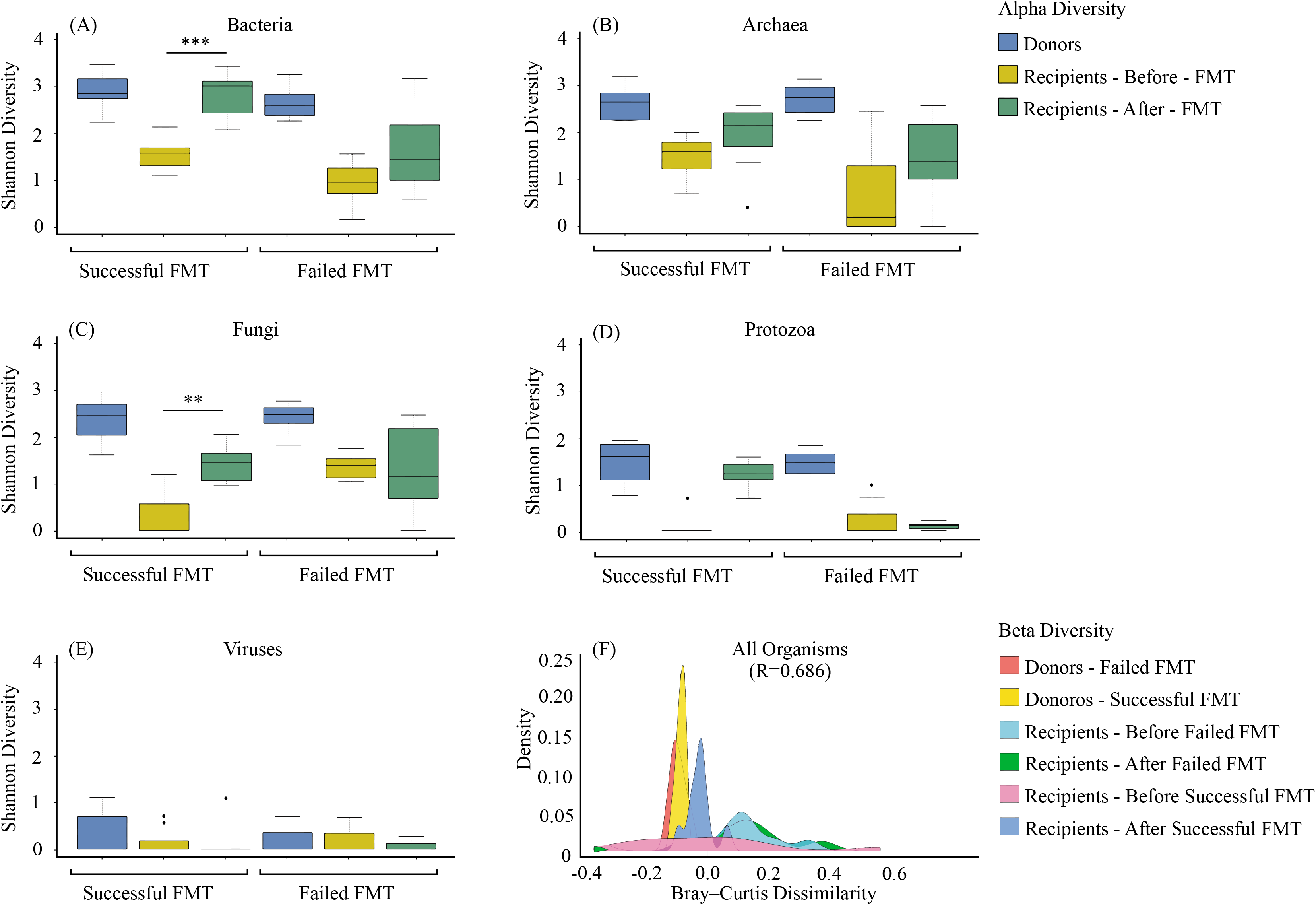
Gut microbial diversity of FMT donors and recipients. The α-diversity (Shannon index) of (A) bacteria, (B) archaea, (C) fungi, (D) protozoa, and (E) viruses of donors, recipients pre- and post-FMT for successful and failed FMT outcomes of rCDI patients. Significant differences were determined using the Kruskal-Wallis and Wilcoxon signed-rank tests for unpaired and paired (i.e. when analysing pre- and post-FMT of recipients) samples, respectively, followed by Bonferroni post-hoc correction. Adjusted p-values were defined at *p<0.05, **p<0.01, ***p<0.001, and ****p<0.0001. The Beta diversity was calculated for all microorganisms (F) using the Bray-Curtis dissimilarity and analyzed using ANOSIM.

The microbial composition of donors, recipients pre-FMT, and recipients post-FMT also differed from each other, as measured by the Bray-Curtis based density plots (ANOSIM, R=0.686, p=0.001) (Fig. 1F). Specifically, pre-FMT, the microbial community of recipients deviated from those of healthy individuals based on Bray-Curtis dissimilarities (ANOSIM, R=0.920) while the microbial community structure of successful and failed donors overlapped (ANOSIM, R=0.648) (Fig. 1F). After successful FMT, the recipients’ microbiome composition resembled the donors (ANOSIM, R=0.595). However, the composition of failed FMT recipients pre and post-FMT overlapped (ANOSIM, R= 0.719) (Fig. 1F).

Since the restoration of bile acid metabolism and SCFA production is reported to account for FMT efficacy, we then extracted bacterial genera associated with these functions (15, 32), as well as sporulating communities (33). Our results showed that successful and failed FMT donors did not significantly differ in their alpha diversity for bile acid metabolizers, SCFA producers, and sporulators (Fig. 2A, B, C). However, a significant increase in the alpha diversity was observed in successful recipients post-FMT for bile acid metabolizers (Wilcoxon signed-rank test, p=0.003), SCFA producers was observed after successful FMT (Wilcoxon signed-rank test, p= 0.014), and sporulators (Wilcoxon signed-rank test, p=0.015) (Fig. 2A, B, C). The density plot for Bray-Curtis dissimilarity showed that the failed and successful FMT donors were not significantly different in their SCFA producing and sporulating communities (ANOSIM, R=0.524 and ANOSIM, R=0.524) (Fig. 2E, F), but differed in their bile acid metabolizing community structures (ANOSIM, R=0.828) (Fig. 2D). Specifically, the relative abundance of bacterial bile acid metabolizers including *Lactobacillus* (associated with deconjugation and esterification of bile salts), *Fusobacterium* (associated with desulfation of bile salts), *Pseudomonas* (desulfation of bile salts), and *Escherichia* (oxidation and epimerization of bile salts) were significantly lower in unsuccessful donor samples (Fig. 3A). Interestingly, intra-variability within donors pertaining to the abundance of bacterial bile acid metabolizers can also be observed (Fig. 3A), which shows that donor composition can vary over time and affect FMT outcome. The successful FMT recipients pre- and post-FMT also differed in their community structure for bile acid metabolizers (ANOSIM, R=0.670), SCFA producers (ANOSIM, R=0.759), and sproulators (ANOSIM, R=0.872) while the failed FMT recipients pre and post-FMT overlapped for the community structure of genera associated with these functions (ANOSIM, R=0.091, R=0.117, and R=0.134, respectively) (Fig. 2D, E, F). Interestingly, the relative abundance of all investigated bacterial bile acid producers was decreased in the failed FMT recipients post-FMT compared to successful FMT recipients (Fig. 3C). Therefore, successful FMT is associated with the colonization of bile acid metabolizing bacteria in the recipients.

**Fig 2.**
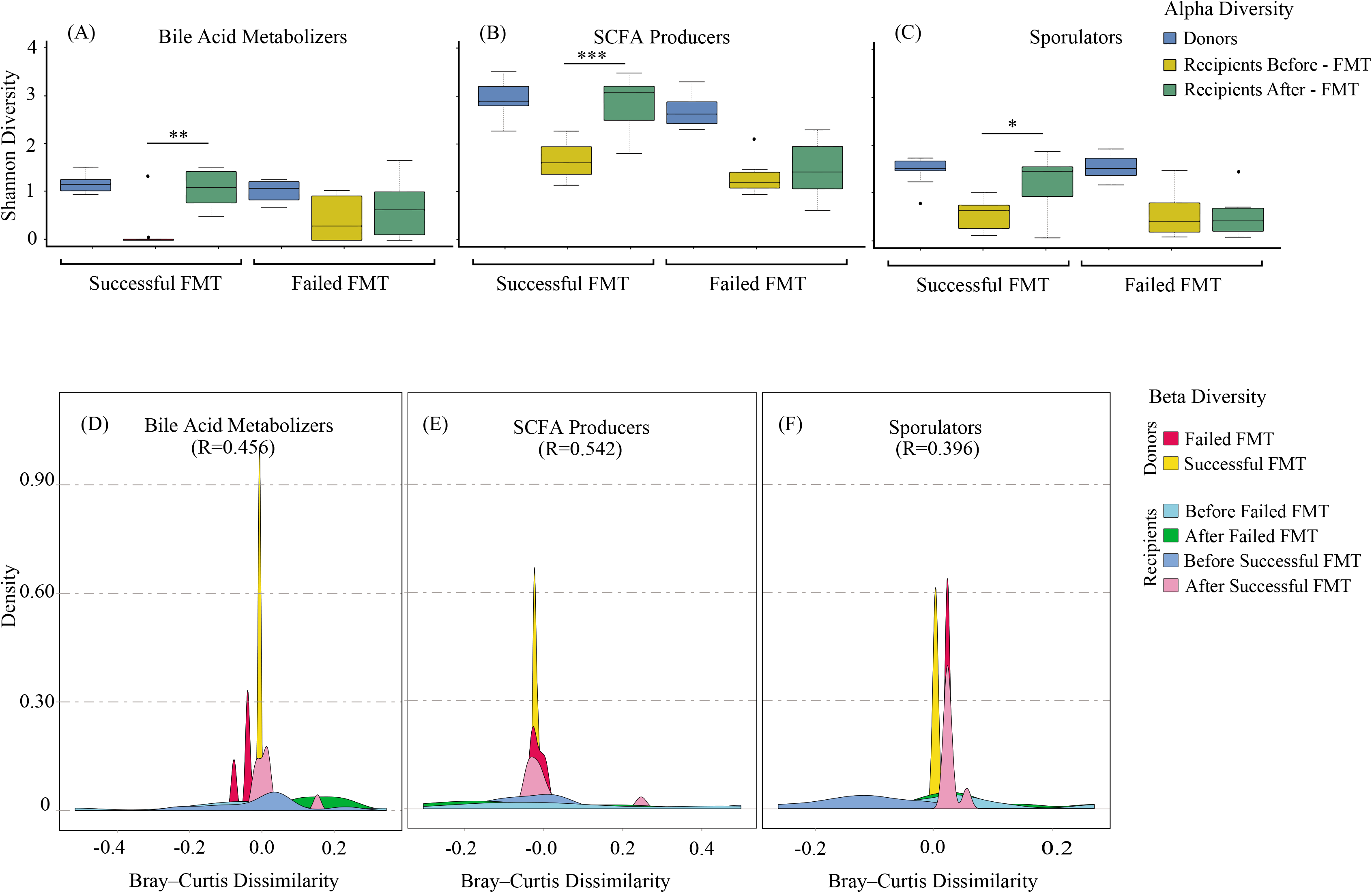
Gut microbial diversity of bile acid metabolizers, SCFA producers, and sporulators. The α-diversity (Shannon index) of (A) bile acid metabolizers, (B) SCFA producers, and (C) sporulating bacteria of donors, recipients pre- and post-FMT for successful and failed FMT outcomes. Significant differences were determined using the Kruskal-Wallis and Wilcoxon signed-rank tests for unpaired and paired samples, respectively, followed by the Bonferroni post-hoc correction. Adjusted p-values were defined at *p<0.05, **p<0.01, ***p<0.001, and ****p<0.0001. The Beta diversity was also calculated using the Bray-Curtis distance-based density plots (D, E, F) and analyzed using ANOSIM.

**Fig 3.**
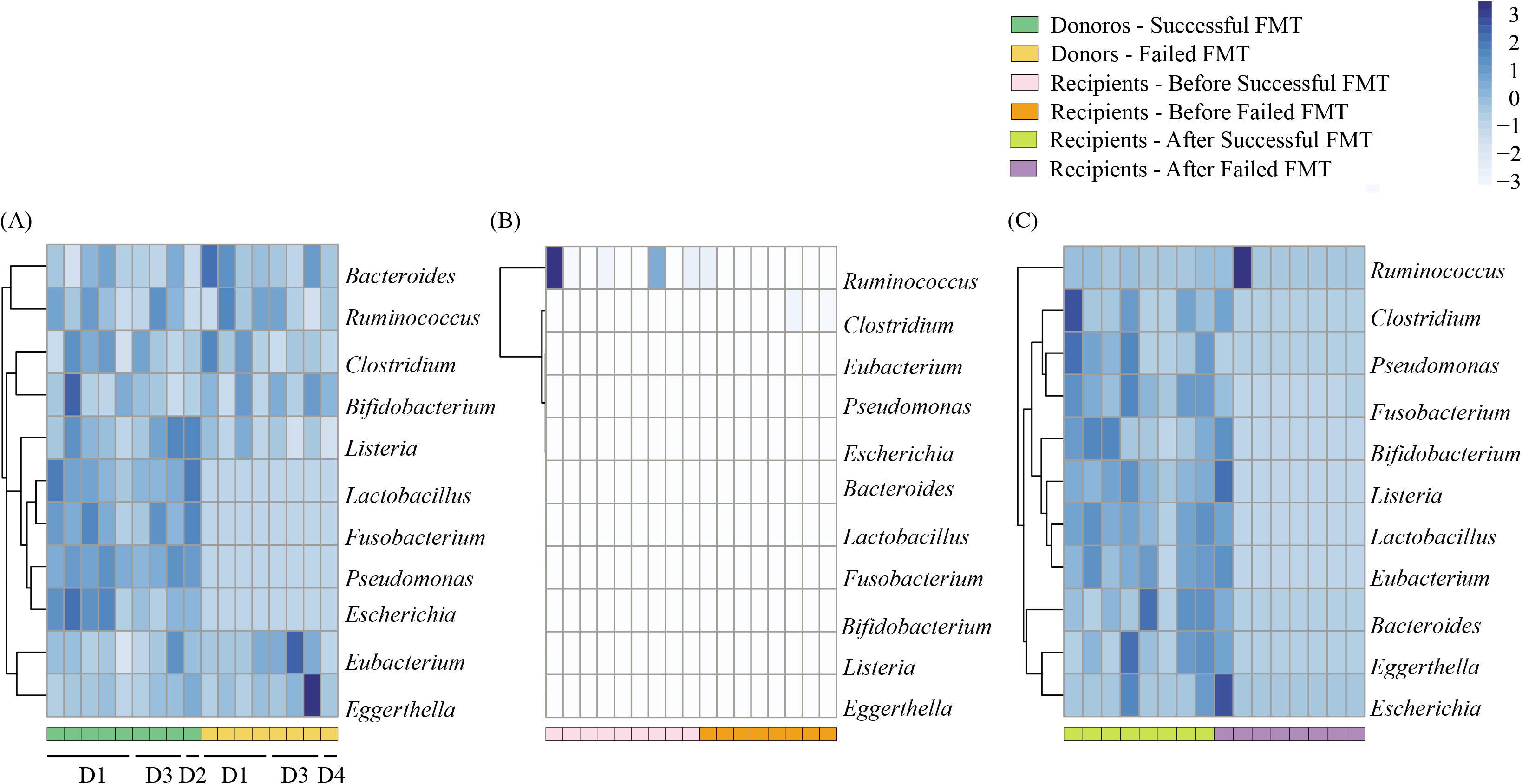
Bile acid metabolizers of FMT donors and recipients. Heatmap of bile acid metabolizers of (A) donors (D1-D4), (B) recipients pre-FMT, and (C) recipients post-FMT for successful and failed FMT outcomes. The dendrogram shows clustering based on the relative abundances. The heatmap color (white to dark blue, corresponding to low to high) represents the row z-score of the mean relative abundance values.

In addition, we also investigated whether microbial community compositions of donor and recipients before FMT can predict the treatment outcomes. The top 20 features from PCA analysis were selected and employed in the subsequent training process of a classification model, using samples from both donor and recipient pre-FMT at the genus level. Using LOO cross validation, the prediction model was significant (p=0.0099) (Fig. 4A), with the most important genera being *Desulfovibrio, Filifactor, Bacillus, Yarrowia, Odoribacter, Wigglesworthia, Oscillibacter, Intestinimonas*, and *Clostridiodes* (Fig. 4B). Furthermore, in order to visualize the impact of the top features on FMT efficiency, the relative abundance of such features was plotted for donor and recipient samples pre- and post-FMT (Fig. 4C). Interestingly, the fungal genus of *Yarrowia*, as well as bacterial genus of *Wigglesworthia*, were significantly higher in pre-FMT failed recipients than pre-FMT successful recipients (Fig. 4C, Kruskal-Wallis, p=0.001 and p=0.002, respectively). The donor samples that contributed to a successful FMT outcome had a higher abundance of *Clostridiodes* (p=0.002), *Desulfovibrio* (p=0.004), *Odoribacter* (p=0.002), and *Oscillibacter* (p=0.003) compared to failed FMT donors (Fig. 4C), and intra-variability in the relative abundances of these genera for each donor was observed (Fig. S1) when comparing successful and failed samples. It is important to note that, these genera were not detected in successful recipients post-FMT (Fig. S1 and S2), indicating that long-term colonization of these genera in recipients may not be critical for FMT success.

**Fig 4.**
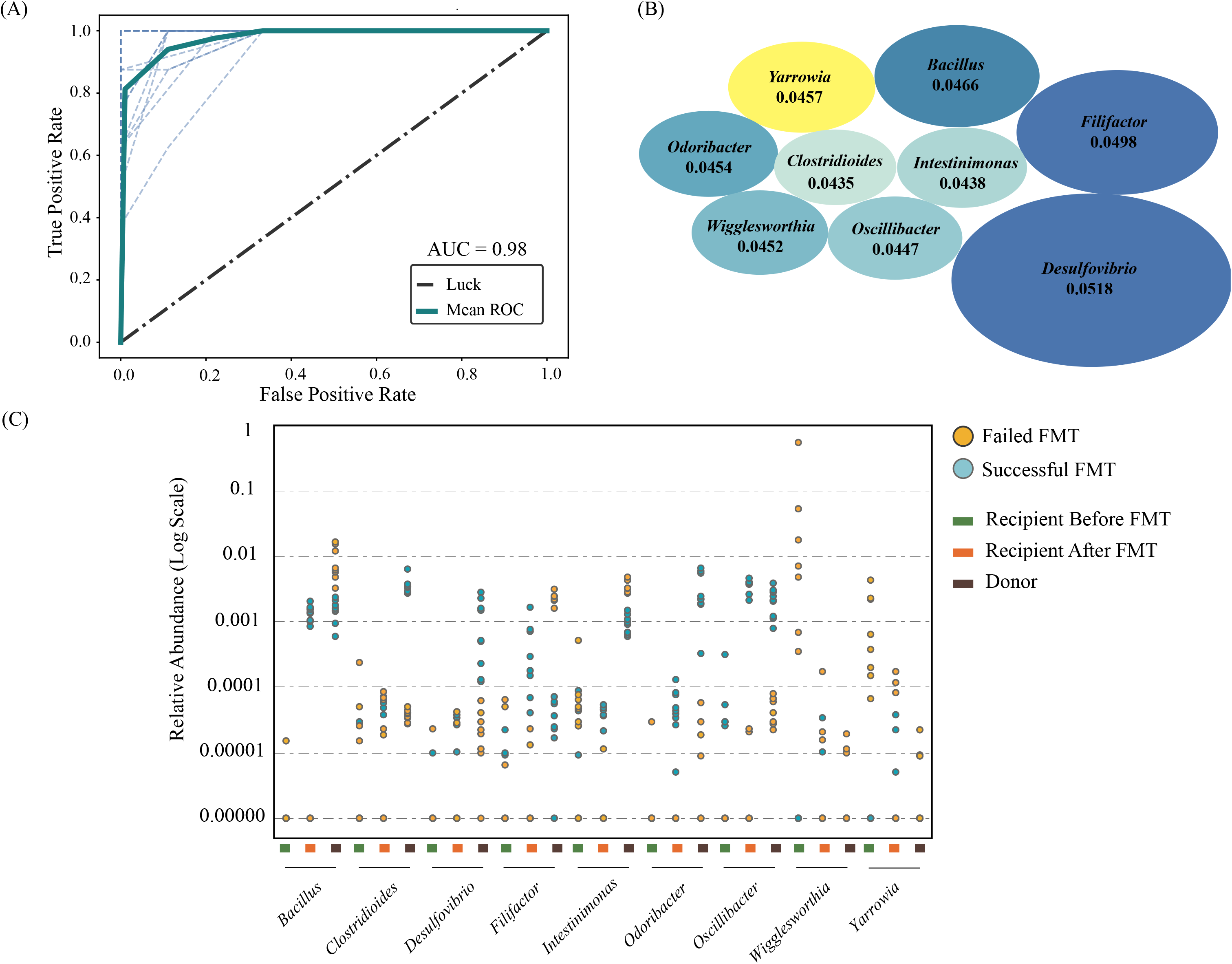
Machine learning model predicting FMT outcome. The random forest model utilized led to (A) the ROC curve displaying an AUC of 98%, (B) the top 9 most important microbes (Blue ellipses represent bacteria; Yellow ellipse represent fungi) involved in FMT success prediction where the size of the ellipses represents the feature importance magnitudes (average Mean Decrease in Impurity), and (C) the relative abundance (shown on logarithmic scale) for the top 9 features of donor, and recipient samples pre- and post-FMT.

When we evaluated our model’s performance against an independent dataset (34, 35), interestingly the model identified similar top features including *Odoribacter* and *Clostrioides*; albeit with no statistically significant discriminatory powers. This was expected due to technical variation between studies which overshadowed the biological variation as well as the lack of full consistency between the two studies rooted in the difference in the average age or ethnicity of the two cohorts.

## Discussion

Despite long-term stability and plasticity of healthy and low to moderately disturbed gut systems (36), severely damaged gut ecosystems are not self-renewing; therefore, FMT can help with restoring damaged systems through (a) the recreation of the original ecosystem (e.g., by autologous FMT) or (b) the construction of an entirely new and alternative ecosystem (e.g., by allogeneic FMT). In our study, we showed that the success of gut ecological recovery through FMT is dependent on several factors, including the donor gut microbiome (the presence of specific bacteria) as well as the pre-FMT recipient gut community structures and recovering habitat (the absence of specific bacteria and fungi) (Fig. 4). In addition, short-term fluctuations in the gut microbiome of both donors and recipients have profound implications in FMT success by producing temporary changes or loss of function (see supplemental materials Fig. S1 and S2; Fig. 3). Therefore, the notion of the “super-donor” is oversimplified due to the observed short-term fluctuations, and a recipient’s microbiota may be just as important to consider when predicting treatment outcomes, especially in other dysbiotic conditions such as ulcerative colitis.

Our results also showed that a trans-kingdom interaction between bacteria and fungi may be important to consider in FMT outcomes. Considering ecological theories on community construction and recovery after disturbance, we hypothesize that the first step of a successful FMT is the colonization of “nexus species” including members of *Desulfovibrio, Odoribacter, Oscillibacter*, and *Clostridioides* genera, as identified in two independent datasets (Fig. 5). These are transient in the community development, but are ecosystem engineers that determine secondary succession trajectories of the ecosystem (Fig. S1 and S2). For example, *Odoribacter* is a known SCFA producer (37). Thus, its presence in the donor and the initial transfer to recipients may contribute to decreased inflammation (38). In addition, the class Clostridia includes many endospore-forming organisms that have the capacity to produce SCFAs (39, 40), which can induce T regulatory cells and associated anti-inflammatory cytokines (17). Following a successful repair, the secondary succession of endogenous or exogenous bile acid metabolizers can restore microbial diversity (lost commensals) and a variety of ecosystem functions (41). Namely, when bile acid metabolizers colonize the repaired gut ecosystem, secondary bile acid concentrations, as pleiotropic signaling molecules in the gut, liver, and systemic circulation, increases (42). This process entails the germination of endogenous or exogenous sporulators such as *Clostridia* and other putative endospore formers, which are considered stress-resistant and are particularly adaptive to cross-host dissemination (19, 43). Aligned with the above hypothesized mechanism, donors that led to a failed FMT had reduced *Fusobacterium* and *Pseudomonas* genera, which are both capable of desulfating primary bile acids. When these genera exist, sulfation can reduce primary bile acid toxicity and increase secondary bile acid excretion via urine and feces (44). This reduced desulfation capacity in failed donor samples further perpetuates the already existing disturbed bile acid pool and inhibits successful secondary colonization for functional ecosystem restoration. Moreover, bacterial genera, which can dehydroxylate primary bile acids into secondary bile acids, are also known to produce SCFAs (38). These gut microbiota associated metabolites, especially butyrate, are a main source of energy for colonocytes and can activate G-protein coupled receptors that regulate intestinal motility and inflammation (38, 45). Lack of such genera in donor samples may diminish the therapeutic potential of FMT.

**Fig 5.**
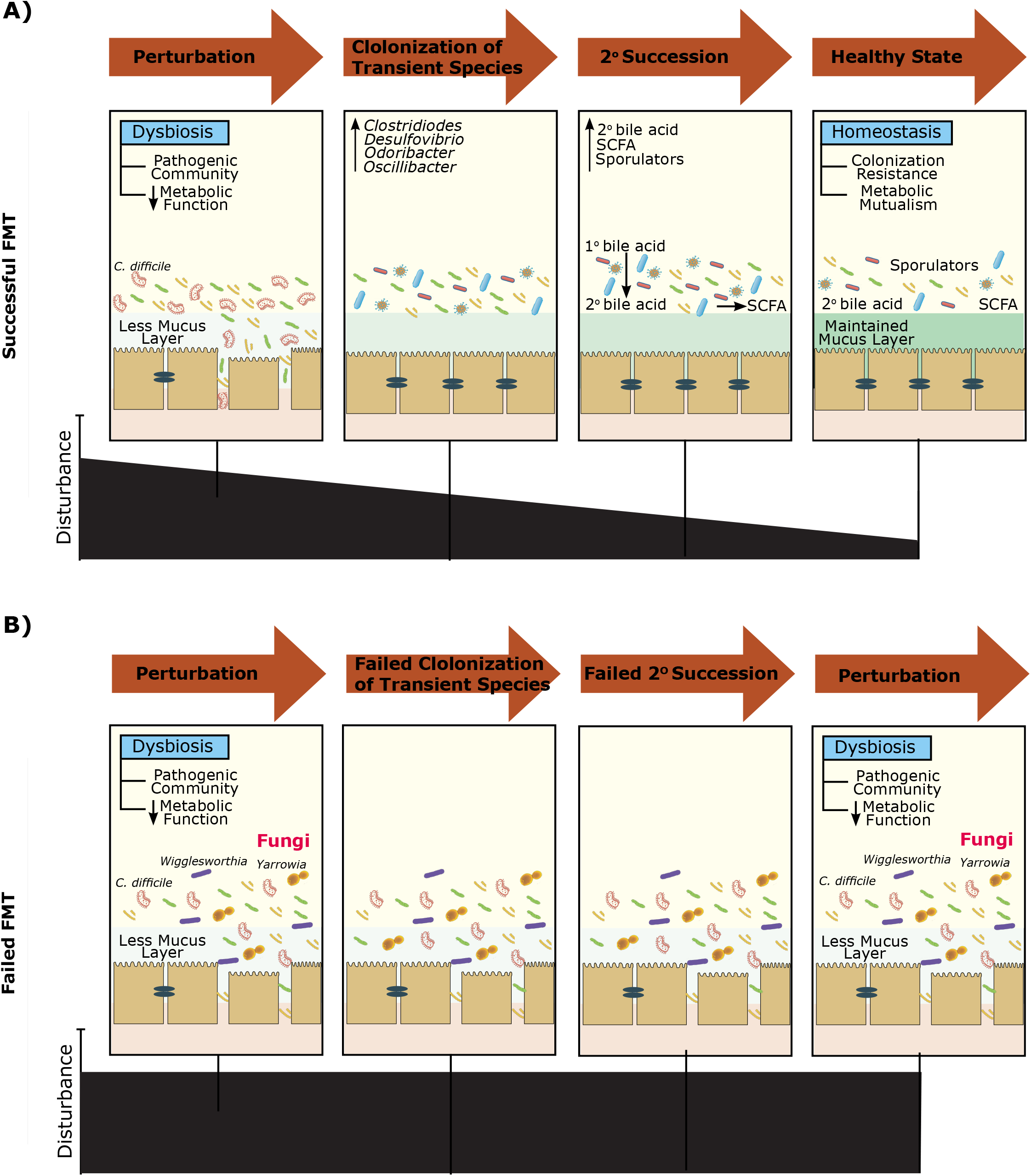
Multifaceted mechanisms affecting FMT treatment outcome. FMT treatment outcome of (A) successful FMT recipients, and (B) failed FMT recipients. A successful treatment outcome includes the repair of the disturbed gut microbial ecosystem by transient colonization of nexus species followed by secondary succession of bile acid metabolizers, sporulators, and short chain fatty acid producers. A failed treatment outcome may be due to the presence of fungal and bacteria genera including *Yarrowia* and *Wigglesworthia* in recipients, minimizing the establishment of repair or successful secondary colonization for functional ecosystem restoration.

However, interestingly, the presence of the *Yarrowia* and *Wigglesworthia* genera in pre-FMT recipients can act as a barrier for the establishment of repair or successful secondary colonization for functional ecosystem restoration (Fig. 4C). This can be due to nutrient cycling and carbon uptake elevation by fungal activity. Moreover, *Yarrowia lipolytica* has been vastly studied as a non-conventional yeast species capable of synthesizing a group of metabolites, in particular lipases and other hydrolytic enzymes (46). These opportunistic fungal pathogens can cause infections in immunocompromised and critically ill patients (47–49). To overcome this challenge, treatment targeted at these fungal elements prior to FMT may potentially enhance treatment efficacy.

In summary, the findings presented herein shed light on potential interlocked mechanisms underlying FMT treatment success, beyond the bacterial community effect. Additionally, FMT is not a ‘one stool fits all’ approach and a more personalized treatment with the inclusion of both donor and patient variables should be taken into consideration to maximize the chance of FMT success. Further knowledge of these factors and mechanisms would be required to optimize FMT treatment for rCDI and possibly other dysbiotic-related diseases. However, the results should not yet be generalized to other patient populations with different demographic characteristics, since our study cohort was small with 88% Caucasian. This signifies the need for larger cohort studies that include patients with diverse demographic characteristics.

## Materials and Methods

### Study design and sample collection

Seventeen adult male and female patients who received FMT for rCDI at the University of Alberta Hospital in Edmonton, Alberta, Canada, between October 2012 and November 2014 were included in this study (31). Criteria for receiving FMT were 1) at least 2 recurrent episodes of mild to moderate CDI, or 2) at least 1 recurrent episode of CDI requiring hospitalization. This study was approved by the University of Alberta Health Research Ethics Board, and all participants provided written informed consent. All participants received FMT by colonoscopy, with stool samples from unrelated donors registered with the Edmonton FMT program. Donor selection criteria and screening process have been described previously (50). After a failed FMT, each patient received FMT from the same donor or a different donor, depending on donor availability. Patients discontinued antibiotics for CDI 24 hours prior to FMT and took 4 L of polyethylene glycol-based bowel preparation (GoLYTELY) one day prior to FMT. Stool samples were collected from donors and patients one week prior to FMT (pre-FMT) as well as from patients one week following FMT (post-FMT). Figure 6 shows the number of donors and recipients, as well as the FMT treatment outcomes. It’s important to note that although some donors had provided multiple stool samples, these samples were provided at different time points (minimum of a one-week gap), which were then administered to the recipients (Fig. 6). It has been perceived that the autocorrelation between microbiomes of stool samples of a given donor normally diminishes between 3-5 days (51).

**Fig 6.**
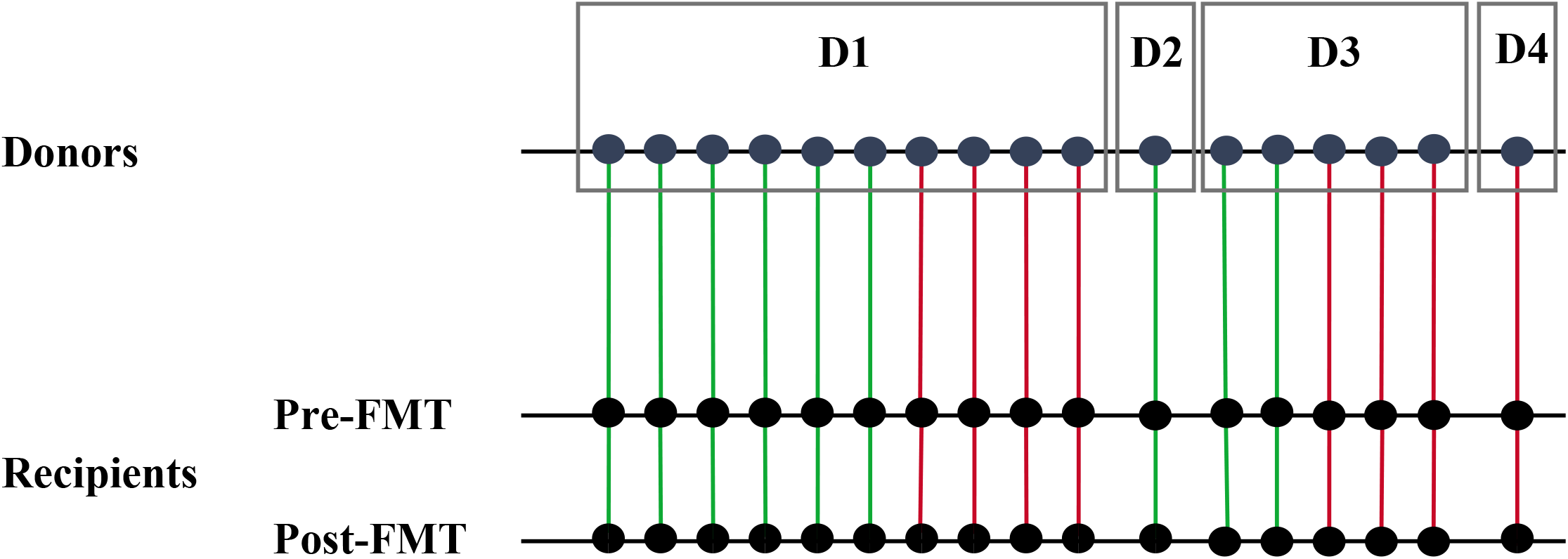
The experimental data structure of stool samples collected. Samples were collected from 4 donors (D1 to D4) and 17 patients one week prior to FMT (pre-FMT) and patients one week following FMT (post-FMT). For D1 and D3, multiple independent samples were taken for different patients. Green line indicates a successful FMT outcome and a red line indicates a failed FMT outcome.

### Metagenomic data collection

DNA from stool samples were extracted using the Qiagen QIAamp DNA stool kit. Shotgun sequencing for metagenomics was applied using the Nextera XT DNA Sample Preparation Kit, and Illumina MiSeq platform was performed as previously described (31). Host DNA was detected and reads were removed by mapping with the GEM program to the human genome with inclusive parameters (52). A custom Kraken database was built of whole genomes of bacteria, viruses, archaea, eukaryote, and viroids (53). The Bayesian Reestimation of Abundance with KrakEN (Bracken) algorithm was used (kmer length of 30 and read length of 100 bp) to compute the abundance of species in DNA sequences originating from each metagenomic sample (54). Singletons, as well as those taxa occurring in only one or two samples, were removed and abundances of different microbial genera were obtained by collapsing detected taxonomies to the genus level and summing features within the same genera. Subsequently, taxa abundances were normalized by the total number of reads sequenced in each sample.

### Statistical analysis

The α-diversity (Shannon diversity index) of successful and failed FMT samples were compared for all organisms using the R package Vegan (55). In addition, α-diversity of bacteria related to bile acid metabolizers (32). SCFA producing genera (15) and sporulators (33) were compared for successful and failed FMT samples. Significant differences in α-diversity were determined using the non-parametric Kruskal-Wallis and Wilcoxon signed-rank test for unpaired and paired samples (pre- and post-FMT samples of recipients), respectively, using Bonferroni correction to adjust the probability. Differences among community structures across samples (β-diversity) were calculated using the Bray-Curtis dissimilarity metric using the R package Vegan and visualized via density plots using custom python scripts (55). Significant differences in β-diversity across donors and recipients were evaluated using analysis of similarities (ANOSIM) (56). Heatmap clustering graphs were constructed using the R pheatmap package to visualize the relative abundance of major bile acid producers in donors and recipients before and after FMT (57).

To test whether donor and recipient microbial composition can predict FMT outcome, we trained a Random Forest (RF) model on pre-treatment samples of both donors and recipients at the genus leve (58). The microbial taxa of both donors and recipients constitute the feature space of the model and the following steps were performed using the Python library, Scikit-learn (59). As the features’ count outnumbers that of the test samples, a dimensionality reduction method was implemented so that the trained model avoids overfitting and generalizes better on the test data (60). Thus, the Principal Component Analysis (PCA) was used to exploit the features which describe the principal components the most. The top 20 features from this analysis were selected to be employed in the training process of the RF model. In order to assess how well the trained classifier generalizes in case of unseen data, the Leave One Out (LOO) cross-validation method was employed. In this method, each data point was used once as a test data, while the classifier was trained on the remaining data points. Subsequently, the cross-validation error value was calculated by averaging all the measured test errors. For each LOO data subset, the Receiver Operating Characteristic (ROC) curve was plotted. Next, the RF classifiers with the highest validation scores were compared by implementing a statistical significance test. Herein, McNemar’s test was used to determine the statistical significance of the difference between the predictive performance of the top RF candidates (61). The RF model identified to be the most precise was then employed to find the most important features in the FMT treatment outcome task. After running the model 100 times, the average Mean Decrease in Impurity (MDI) of the most important features were also calculated (62). Subsequently, the Kruskal-Wallis test with the Bonferroni correction to adjust the probability was utilized to compare the relative abundance of the top important features across the samples. Lastly, in an attempt to evaluate our model’s performance and its generalizability another independent dataset was used (34, 35). This dataset consisted of DNA extracted from 5 fecal samples from 3 donors, and 5 fecal samples from each of 10 FMT recipients: collected at day 0 (pre-FMT) and days 2, 14, 42 and 84 after FMT.

## Supplemental Material

**Fig. S1** The intra-variability of relative abundance of the top 9 features of donor 1 (D1) samples and their corresponding recipients, pre- and post-FMT leading to successful (A and B) and failed (C and D) FMT outcomes.

**Fig. S2** The intra-variability of relative abundance of the top 9 features of donor 3 (D3) samples and their corresponding recipients, pre- and post-FMT leading to successful (A and B) and failed (C and D) FMT outcomes.

## Author Contributions

N.K and S.P wrote the first draft of the manuscript. SP assisted in reviewing literature, guided the analysis, and provided intellectual input in the manuscript. M.R, A.S, and A.N aided with the data analysis. N.K, M.R, A.N., G.K.-S.W, D.K, and S.P reviewed and edited the manuscript. F.M.K and A.H.Z.K contributed to the metagenomic sequence handling and processing. N.K, M.R, A.N., G.K.-S.W, D.K, and S.P actively contributed to the critical discussions. The authors read and approved the final manuscript.

